# DeepScope: Nonintrusive Whole Slide Saliency Annotation and Prediction from Pathologists at the Microscope

**DOI:** 10.1101/097246

**Authors:** S. Joseph Sirintrapun, Hikmat A. Al-Ahmadie, Peter J. Schüffler, Thomas J. Fuchs

## Abstract

Modern digital pathology departments have grown to produce whole-slide image data at petabyte scale, an unprecedented treasure chest for medical machine learning tasks. Unfortunately, most digital slides are not annotated at the image level, hindering large-scale application of supervised learning. Manual labeling is prohibitive, requiring pathologists with decades of training and outstanding clinical service responsibilities. This problem is further aggravated by the United States Food and Drug Administration’s ruling that primary diagnosis must come from a glass slide rather than a digital image. We present the first end-to-end framework to overcome this problem, gathering annotations in a nonintrusive manner during a pathologist’s routine clinical work: (i) microscope-specific 3D-printed commodity camera mounts are used to video record the glass-slide-based clinical diagnosis process; (ii) after routine scanning of the whole slide, the video frames are registered to the digital slide; (iii) motion and observation time are estimated to generate a spatial and temporal saliency map of the whole slide. Demonstrating the utility of these annotations, we train a convolutional neural network that detects diagnosis-relevant salient regions, then report accuracy of 85.15% in bladder and 91.40% in prostate, with 75.00% accuracy when training on prostate but predicting in bladder, despite different pathologists examining the different tissues. When training on one patient but testing on another, AUROC in bladder is 0.7929±0.1109 and in prostate is 0.9568±0.0374. Our tool is available at https://bitbucket.org/aschaumberg/deepscope.

## 1 Introduction

Computational pathology^[10]^ relies on training data annotated by human experts on digital images. However, the bulk of a pathologist’s daily clinical work remains manual on analog light microscopes. A noninterfering system which translates this abundance of expert knowledge at the microscope into labeled digital image data is desired.

Tracking a pathologist’s viewing path along the analyzed tissue slide to detect local image saliency has been previously proposed. These approaches include whole slide images displayed on one or more monitors with an eye-tracker^[5]^, mouse-tracker^[19]^ or viewport-tracker^[21,17]^ – but may suffer confounds including peripheral vision^[16]^, head turning^[1]^, distracting extraneous detail^[2]^, monitor resolution^[20]^, multimonitor curvature^[23]^, and monitor bezel field of view fragmentation^[25]^. Only our approach does not change the pathologist’s medical practice from the microscope. The microscope is a class I device appropriate for primary diagnosis according to the United States Food and Drug Administration, while whole slide imaging devices are class III^[18]^.

In light of the confounds of alternatives, its centuries of use in pathology, and its favorable regulatory position for primary diagnosis, we believe the microscope is the gold standard for measuring image region saliency. Indeed, there is prior work annotating regions of interest at the microscope for cytology technicians to automatically position the slide for a pathologist^[4]^.

We therefore propose a new, noninterfering workflow for automated video-based detection of region saliency using pathologist viewing time at the microscope (Fig 1). Viewing time is known in the psychology literature to measure attention^[13,8]^, and we define saliency as pathologist attention when making a diagnosis. Using a commodity digital camera, rather than a custom embedded eye-tracking device^[7,14]^, we video record the pathologist’s entire field of view at a tandem microscope to obtain slide region viewing times and register these regions to whole slide image scans.

**Figure 1:**
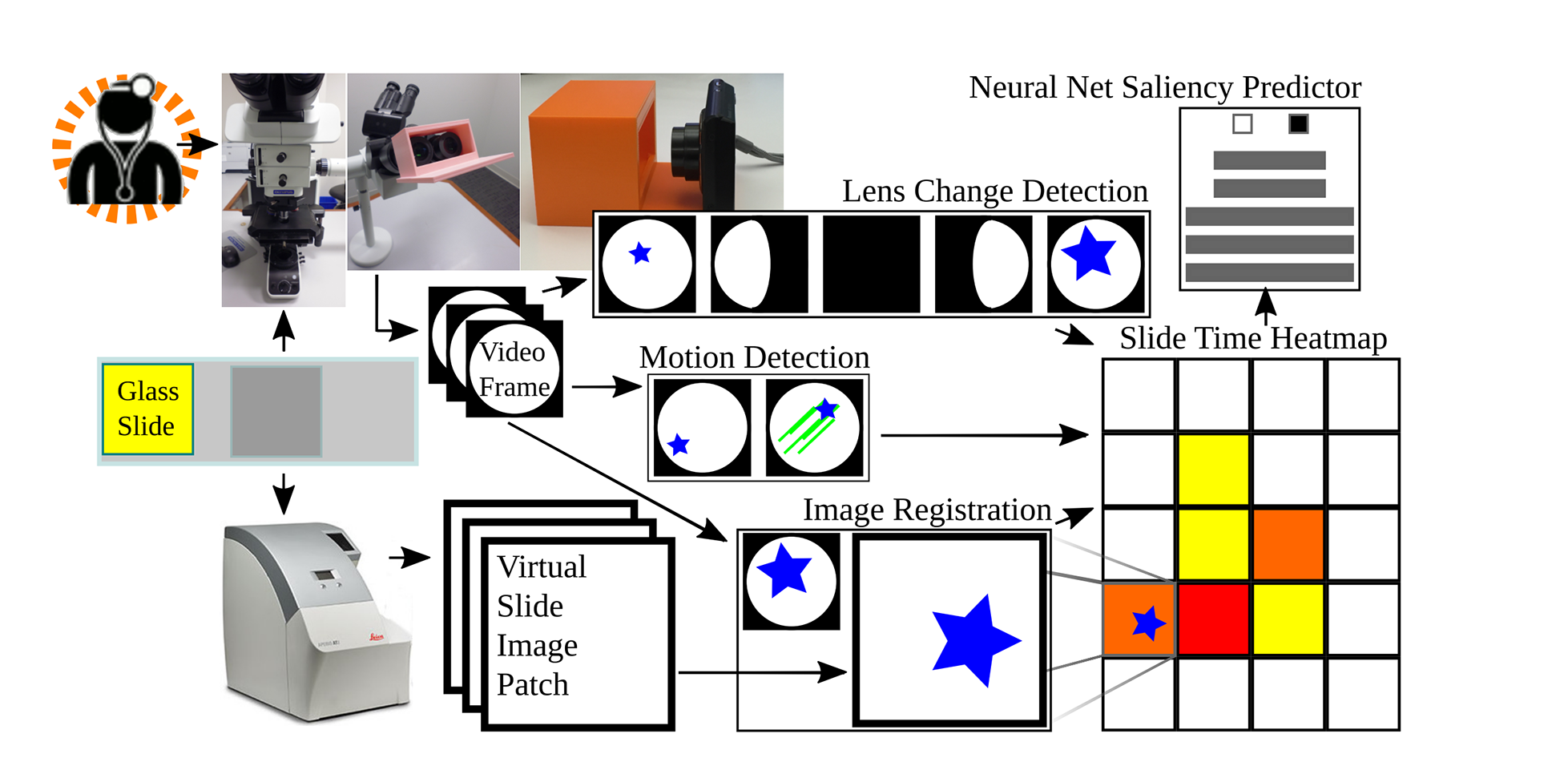
Proposed microscope-based saliency predictor pipeline workflow. The pathology session is recorded, the slide is scanned, the video frames are registered to scan patches. Lens change detection guides registration and viewing time is recorded for periods without motion. A convolutional neural net learns to classify patches as salient (long looks) or not.

Second, we train a convolutional neural network [CNN] on these observation times to predict whether or not a whole slide image region is viewed by a pathologist at the microscope for more than 0.1 seconds. As more videos become available, our CNN predicting image saliency may be further trained and improved, through online learning.

## 2 Methods and Materials

**Pathologists** Pathologists were assistant attending rank with several years experience each. Trainees have different, less efficient, slide viewing strategies^[5,16]^. Region viewing times and path were automatically recorded during a pathologist’s routine slide analysis, without interference.

**Patient slides** Two bladder cancer patients were studied by SJS. Two prostate cancer patients were studied were studied by HAA. One slide per patient was used, for four slides total (Fig 2).

**Figure 2:**
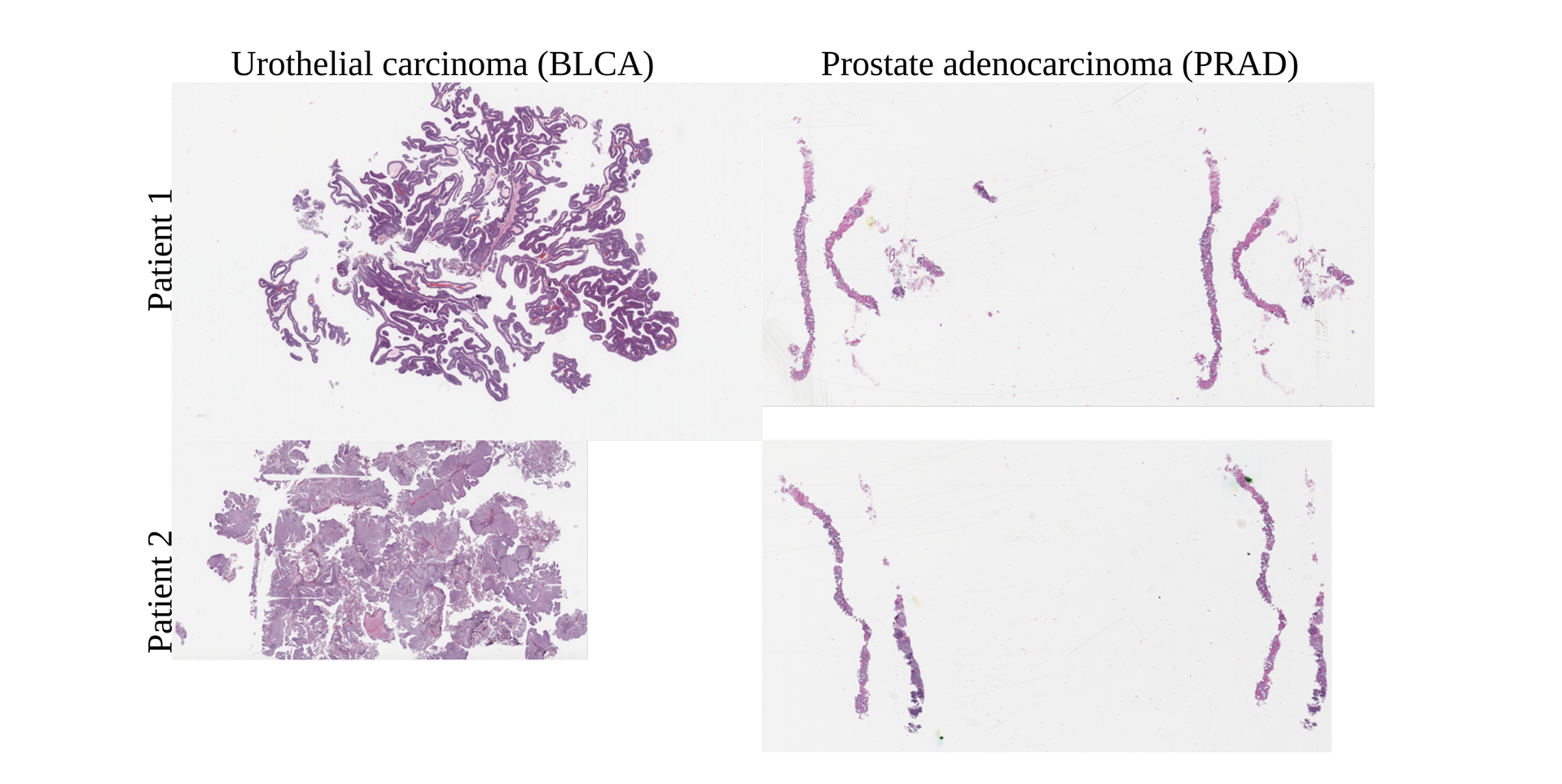
Bladder cancer left, prostate cancer right. Training, validation, testing done on top slides, with additional same-tissue testing on bottom slides. For cross-tissue testing, top slide tested against other top slide. Viewing time heatmap for top left bladder shown in Fig 7. Note how the top bladder has more edges than the more solid bottom bladder, while the prostates have similar tissue texture. We believe this impacts interpatient accuracy, shown in Fig 9.

### Scan preprocessing

Microscope slides, inspected by a pathologist, were scanned at 0.5±0.003 microns per pixel [px], using an Aperio AT2 scanner. The resulting SVS data file consists of multiple levels, where level 0 is not downsampled, level 1 is downsampled by a factor of 4, level 2 by a factor of 16, and level 3 by a factor of 32. From each level, 800x800px patches were extracted via the OpenSlide software library^[11]^. In bladder, adjacent patches in a level overlap at least 50%, to avoid windowing artifacts in registration. In prostate, adjacent patches overlap at least 75%, to best center the pathologist’s field of view on the little tissue in a needle biopsy. Patches evenly cover the entire level without gaps. Scans were either taken before a technician applied marker to the slides, to indicate regions of interest to the pathologist, or after markings were scrubbed from the slide. However, these marks were evident in the pathologist videos discussed in the next section.

### Video acquisition

A Panasonic Lumix DMC-FH10 camera with a 16.1 megapixel charge-coupled device [CCD], capable of 720p motion JPEG video at 30 frames per second, was mounted on a second head of an Olympus BX53F multihead teaching microscope to record the pathologist’s slide inspection. Microscope objective lens magnifications were 4x, 10x, 20x, 40x, and 100x. Eyepiece lens magnifications was 10x. The pathologist was told to ignore the device and person recording video at the microscope during inspection. The mount (Fig 1) for this camera was designed in OpenSCAD and 3D-printed on a MakerBot 2 using polylactic acid [PLA] filament.

### Camera choice

Many expensive microscope-mounted cameras exist, such as the Lumenera INFINITY-HD and Olympus DP27, which have very good picture quality and frame rate. The Lumenera INFINITY-HD is a CMOS camera, not CCD, so slide movement will skew the image rather than blur it, and we did not want to confound image registration or motion detection with rolling shutter skew. Both cameras trim the field of view to a center-most rectangle for viewing on a computer monitor, which is a loss of information, and we instead assign viewing time to the entire pathologist-viewed 800x800px PNG patch from the SVS file representing the whole slide scan image. Both cameras do not have USB or Ethernet ports carrying a video feed accessible as a webcam, for registration to the whole slide scan. The Olympus DP27 may be accessible as a Windows TWAIN device, but we could not make this work in Linux. Finally, the HDMI port on both carries high-quality but encrypted video information that we cannot record, and we did not wish to buy a Hauppauge HDMI recording device, because we had a cheaper commodity camera on hand already. We also considered automated screenshots of the video feed in Aperio ImageScope as displayed on a computer monitor, but we observed a lower frame rate and detecting lens change is complicated because the entire field of view is not available. Recording low-quality video on a commodity camera to a SecureDigital [SD] memory card is inexpensive, captures the entire field of view, and is generally applicable in any hospital. For this pilot study, we used only one camera for video recording, rather than two different microscope cameras, potentially eliminating a confound for how many pixels are moving during rapid short movements of the slide. For 3D printing requisite camera mounts, open source tools are available.

### Video preprocessing and registration

A Debian Linux computer converted individual slide inspection video frames to PNG files using the ffmpeg program. OpenCV software detected slide movement via optical flow^[9]^, comparing the current video frame with the preceding video frame, shown in Fig 3. We defined slide movement to start if 10% or more of pixels in the entire field of view of the camera have a movement vector of at least one, and defined slide movement to stop if 2% or fewer of the pixels in the entire field of view of the camera have a movement vector of at least one. The entire field of view of the camera is 640x480px, a small subset of these capture the circular field of view at the microscope eyepiece, with the remaining pixels being black (Fig 3). The representative frame among consecutive unmoving frames moved the least. The ImageJ^[22]^ SURF^[3]^^1^ and OpenCV software libraries registered each representative to an 800x800px image patch taken from the high-resolution Aperio slide scanner. Each patch aggregated total pathologist time.

**Figure 3:**
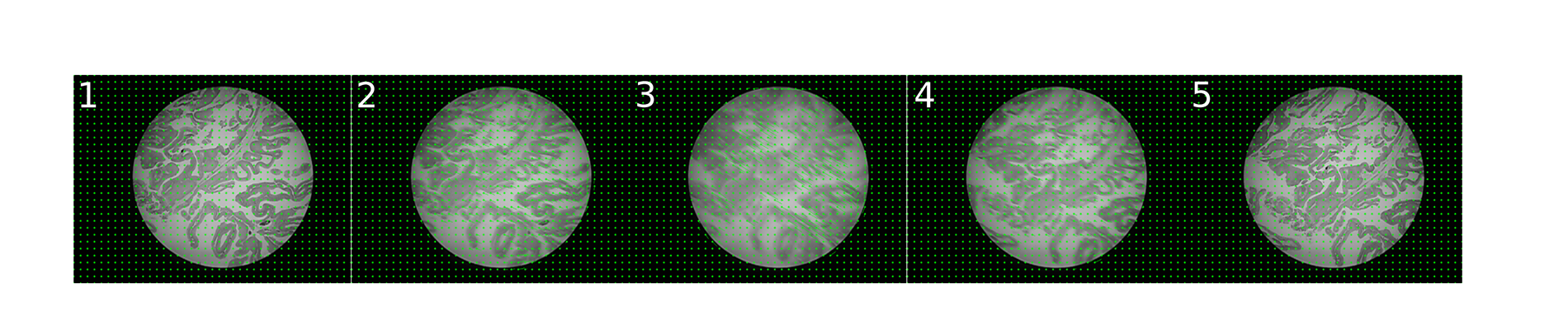
Optical flow, showing pixel movement grid. The frame has few moving pixels before *(left)* and after *(right)* pathologist moves the slide. A pathologist looks at a slide region for the duration of consecutive stationary frames.

The partially automated registration process starts with initial manual registration of a frame, followed by automated registration within the preceding registration’s spatial neighborhood. The OpenCV implementation of random sample consensus [RANSAC] follows a point set registration procedure to calculate a rigid body transformation between the shared SURF interest points in the video frame and an image patch, to find the distance in pixels that the video frame is off-center from the patch, with the least off-center image patch selected as the best registration, because the pathologist’s fovea is in approximately the same place in this video frame and image patch (Fig 4). The ImageJ SURF implementation of interest points shared between two images provides a set of corresponding interest point pairs, one from the frame and one from the patch, enabling point set registration. A final manual curation ensures correctness. This process reduces manual effort in that automatic registrations are rarely far away from the correct registration, and when automatic registrations are incorrect, the manual curation has only a small, local neighborhood to search to make the correction, then automatic registration may proceed from the correction. Fully automated image registration is not part of this study.

**Figure 4:**
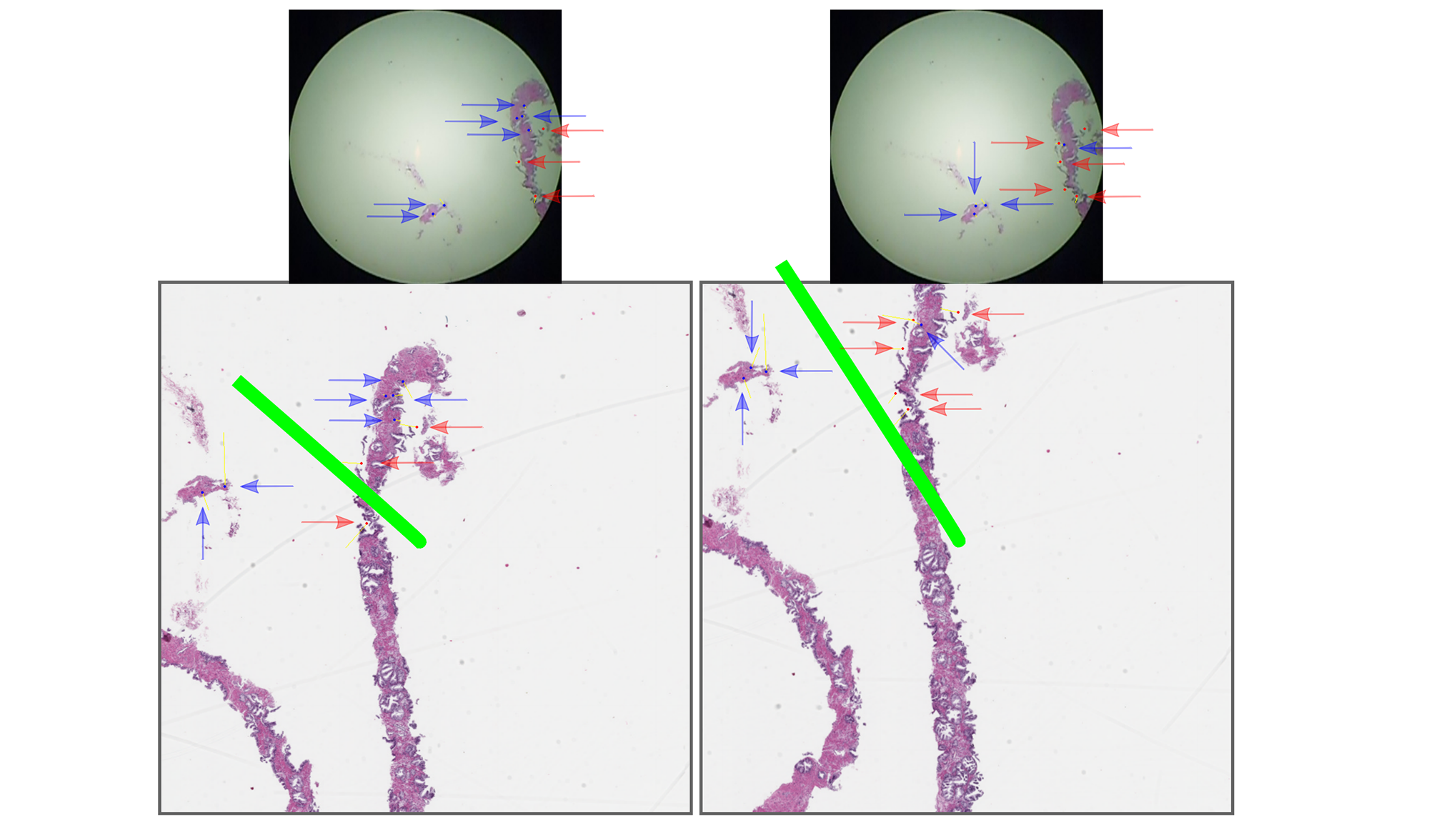
The best image registration for a given video frame (same frame top left and top right) from the commodity camera at the microscope eyepiece compared to two different high-quality patches (bottom left and bottom right) from the whole slide scan image minimizes the length of the green line, which is the distance from the center of the patch to the center of the frame mapped into the patch’s coordinate space.

During slide inspection, the pathologist may switch objective lens magnification. Lens change is detected automatically when the field of view bounding box of nonblack pixels changes size (Fig 5), and is important because SURF is scale-invariant so registrations may otherwise proceed at an unchanged magnification.

**Figure 5:**
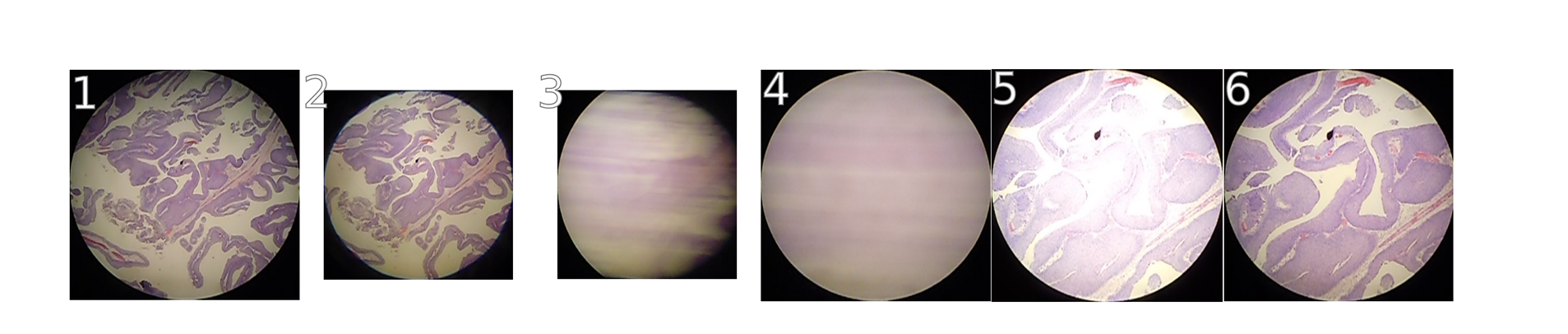
Lens change detection: the normal non-black pixel bounding box is initially 415x415px. A change to 415x282px indicates the pathologist changing the lens, thus changing slide magnification. Note some pixels that may appear black are called non-black due to difficult to perceive noise in the image, which effects calculated bounding box size. All images shown at same scale trimmed to bounding box.

### Deep learning

We used Caffe^[12]^ for deep learning of convolutional features in a binary classification model given the 800x800px image patches labeled with pathologist viewing times in seconds. To adapt for our purpose CaffeNet (Fig 6), which is similar to AlexNet^[15]^, we re-initialized its top layer’s weights after ImageNet^[6]^ pre-training. Two output neurons were connected to the re-initialized layer, then training followed on augmented 800x800px patches for 10,000 iterations in Caffe. In bladder, our model simply predicted whether or not a pathologist viewed an 800x800px patch more than 0.1 sec (30 fps camera). In prostate, due to the higher overlap between adjacent patches and less tissue available, to be salient a patch met at least one of these criteria: (1) viewed more than 0.1 sec, (2) immediately above, below, left, or right of at least two patches viewed more than 0.1 sec, or (3) above, below, left, right, or diagonal from at least three patches viewed more than 0.1 sec such that all three are not on the same side. In this way, image patches highly overlapping in the neighborhood of salient patches were not themselves considered nonsalient if a pathologist happened to jump over them during observation.

**Figure 6:**
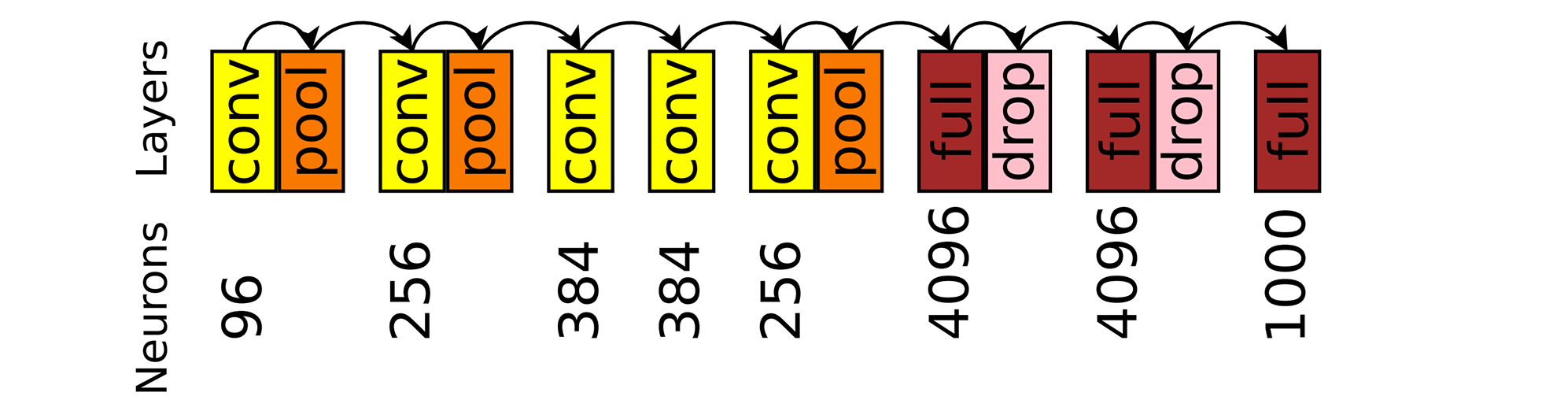
Caffenet neuron counts, convolutional layers, dropout^[24]^ layers, and fully-connected layers.

## 3 Experiments

Urothelial carcinoma (bladder) in Fig 7 was analyzed first, with HAA inspecting at the microscope. Viewed regions at the microscope corresponded to the whole-slide scan SVS file at magnification levels 2 and 1. We restricted our analysis to level 2, having insufficient level 1 data. We split level 2 into three regions: left, center, and right. Due to over 50% overlap among the slide’s total 54 800x800px level 2 patches, we excluded the center region from analysis, but retained the left and right, which did not overlap (Fig 8).

**Figure 7:**
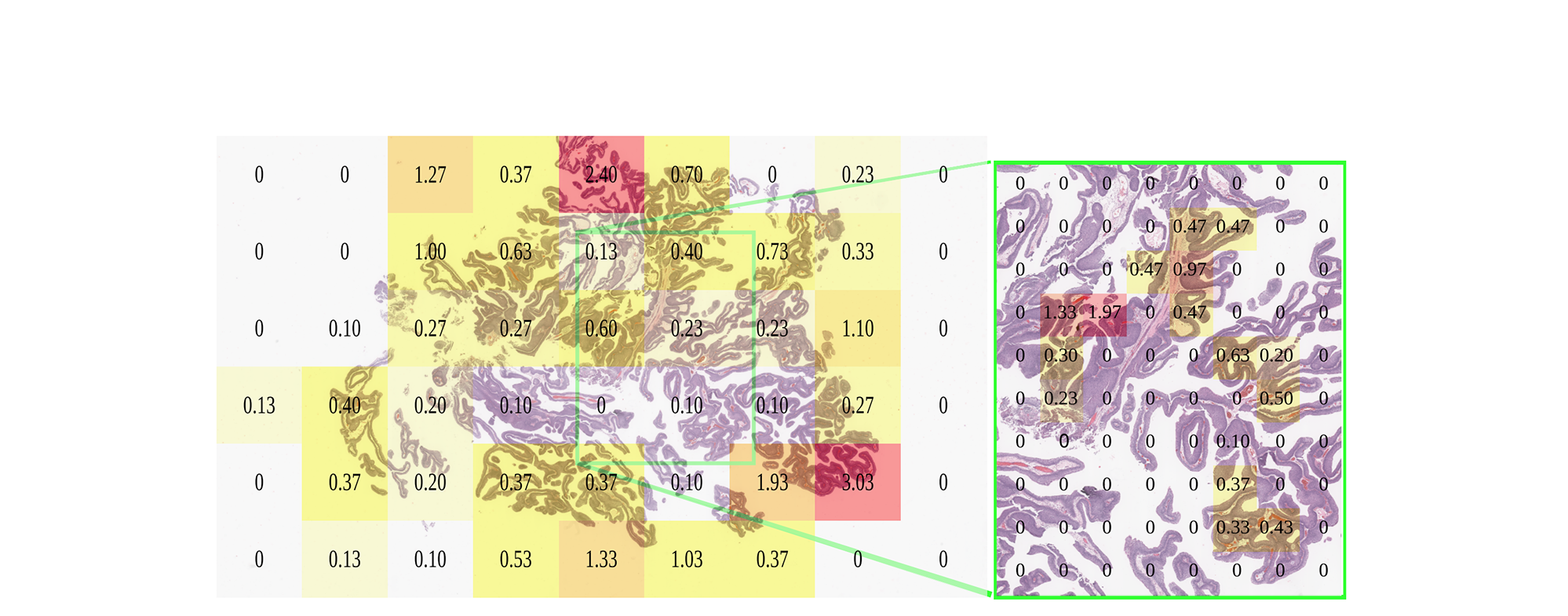
Pathologist viewing times in seconds at the microscope for low (left, 10x, level 2) and high magnification (right, 20x, level 1), registered to the same urothelial carcinoma slide scan.

**Figure 8:**
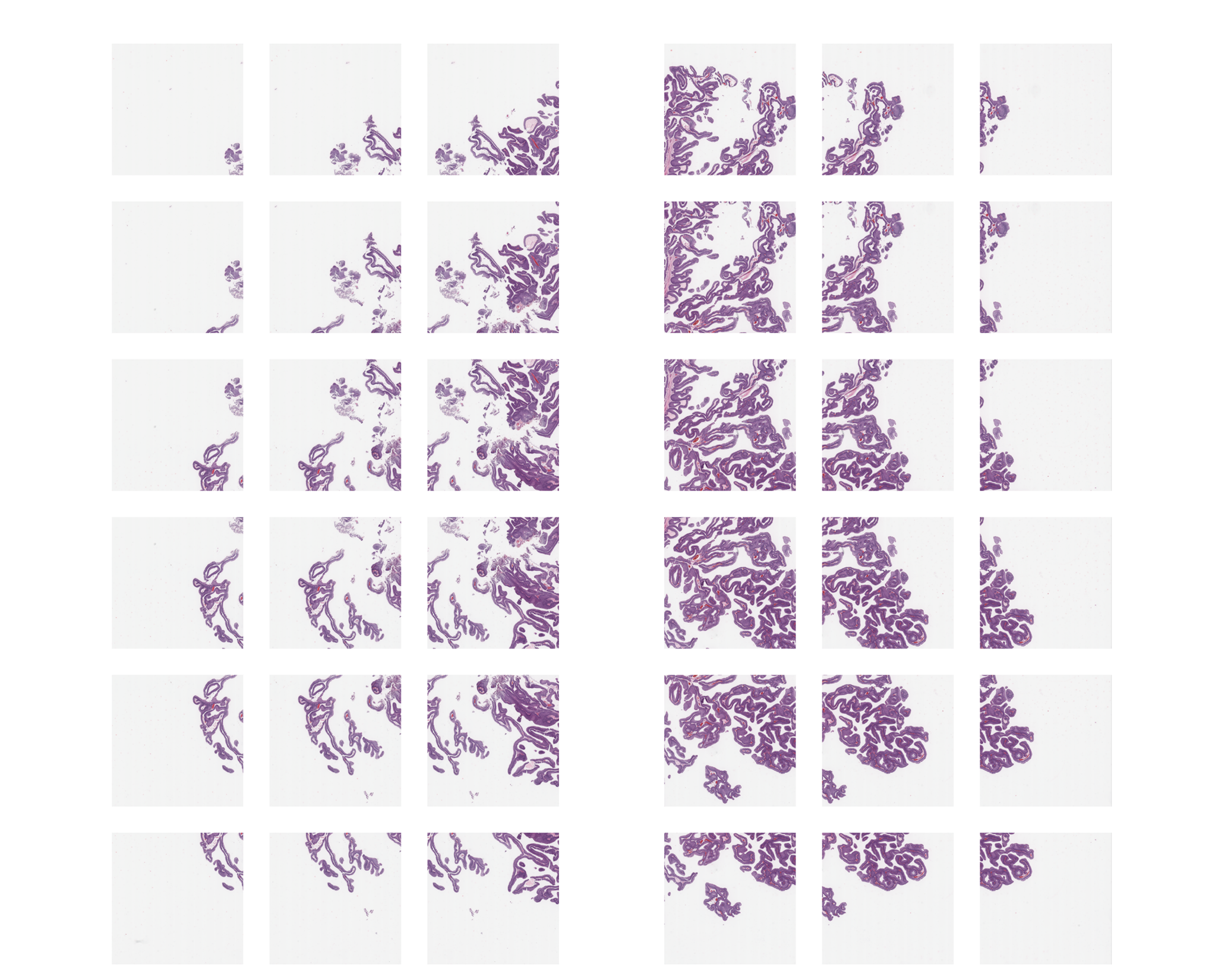
Scaled image patches of left and right sides of bladder patient 1 slide. Middle excluded here and not used in analysis, to isolate left and right sides from each other. Note far left and far right have less tissue, but tissue is present for training. Also note the overlap among patches is evenly distributed and greater than 50%.

In bladder, we considered a negative example to be a patch viewed for 0.1 seconds (3 frames or fewer, 30 fps) or less, and a positive example viewed for more than 0.1 seconds (4 frames or more). This threshold produced 9 positive and 9 negative examples on the left side, and the same number on the right side. We performed three-fold cross validation on the left side (6+ and 6-examples training set, 3+ and 3- examples validation set), then used the model with the highest validation accuracy on the right side to calculate test accuracy, an estimate of generalization error (Fig 9). This cross validation was duplicated ten times on the left side, each time estimating test accuracy, to calculate a confidence interval. We then duplicated this training/validating on the left and testing on the right.

**Figure 9:**
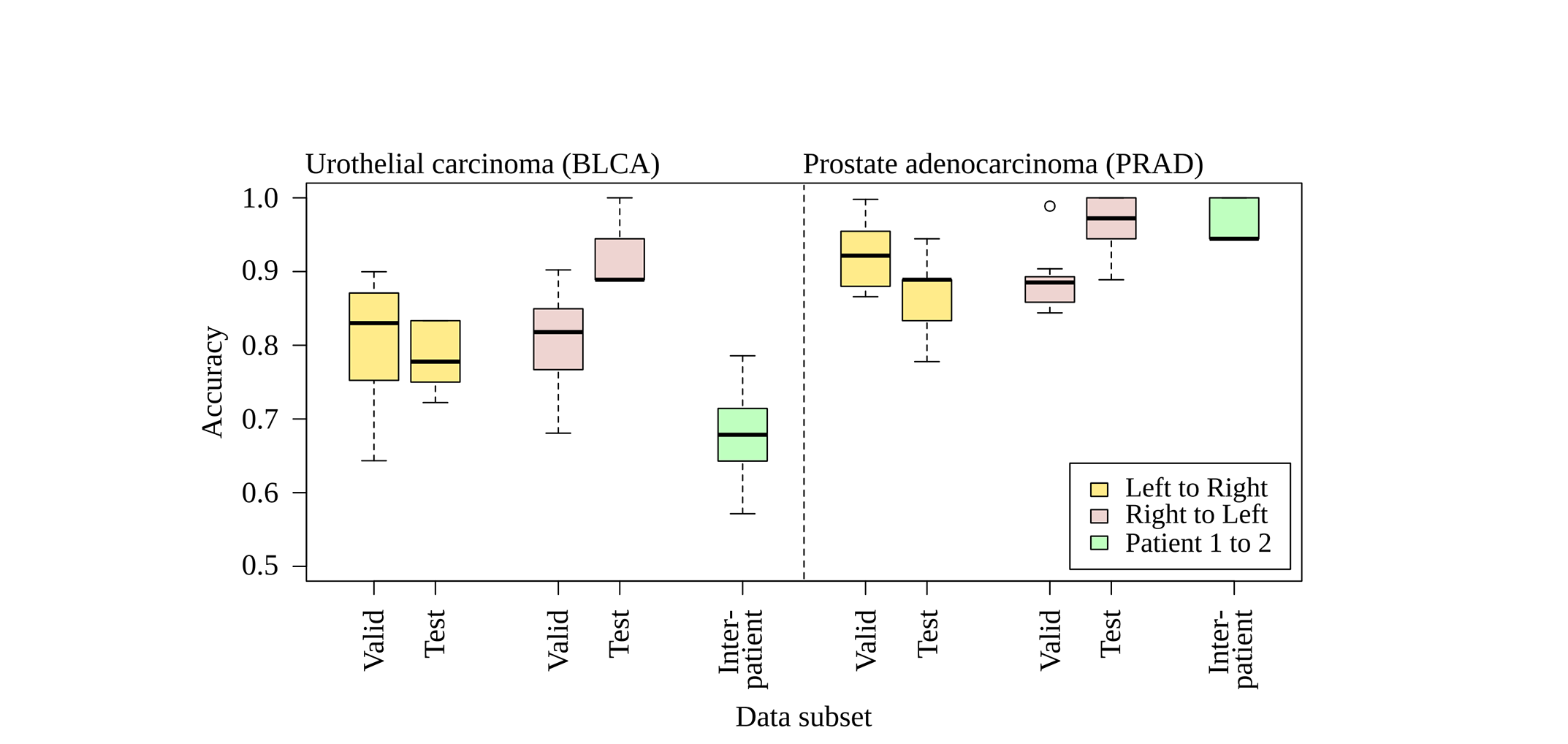
Ten three-fold cross validation trials for bladder [BLCA] and prostate [PRAD], evaluated for intrapatient training/validating on left while testing on the right and vice versa. Each model is evaluated against a different patient (interpatient), slides in Fig 2). The needle for prostate cancer biopsy may standardize the distribution of prostate tissue in the whole slide, maintaining a higher accuracy of the prostate classifier on an interpatient basis than the bladder cancer classifier. The bladder patients are transurethral resections taken by cuts rather than a standard gauge needle.

Training and validation data were augmented. For a 800x800px patch, all 1 degree rotations through 360 degrees were saved, then cropped to the centermost 512x512px, then scaled to 256x256px. Caffe then randomly cropped 256x256px patches to 227x227px for each iteration of CaffeNet learning. No images in the validation set were derived from the training set, and vice versa. A training set is two concatenated folds, with the remaining fold as validation. We used validation accuracy reported by Caffe after training completed, then averaged over all three folds. We did not augment the test set comprising 9 positive and 9 negative examples. In addition to the bladder cancer slide, we analyzed two prostate cancer needle biopsy slides, with SJS inspecting these slides.

## 4 Results

In bladder, when training/validating on the left side and testing on the right, mean test accuracy is 0.781±0.0423 (stdev) with 95% confidence interval [CI] from 0.750 to 0.811 (df=9, Student’s T). When training/validating on the right and testing on the left, mean test accuracy is 0.922±0.0468 with 0.889-0.956 95% CI. Overall mean test accuracy is 85.15%. The left and right test accuracies differ (p=0.000135, Wilcoxon rank-sum, n=20), while validation accuracies do not (p=0.9118, n=20). This may suggest nonhomogenous information content throughout the slide. The pathologist started and ended slide inspection on the right. The second bladder had different morphology and model accuracy reduced to 0.678±0.0772, 0.623-0.734 95% CI. Moreover, the second bladder had only 7 positive examples available, whereas both prostates and the first bladder had at least 9 positive examples available.

For the first prostate slide, training on the left side and testing on the right, we find accuracy 0.867±0.0597, 0.824-0.909 95% CI. Training on the right and testing on left, we find 0.961±0.0457, 0.928-0.994 95% CI. Overall mean test accuracy is 91.40%. Taking the best model learned from this first prostate (right side, test accuracy 100%, 18/18), we tested on the second prostate’s right side (because the left did not have 9 positive training examples) and find 0.967±0.0287, 0.946-0.987 95% CI. We also tested this model on the bladder cancer slide, and find 0.780 accuracy on the left and 0.720 on the right (9+ and 9- training examples each), mean accuracy 75.00%. The best bladder cancer model predicts every patch is not salient in both prostates, presumably because the little tissue in prostate is insufficient for a positive saliency prediction.

Interpatient AUROC for bladder and prostate is shown in Fig 10. In prostate, nine salient and nine nonsalient examples are drawn from the second patient. Average AUROC was calculated from ten such draws, achieving a mean±stdev of 0.9568±0.0374 and 95% CI of 0.9301-0.9835. Over all 17 salient and 13 nonsalient patches used from the second prostate patient, the AUROC is 0.9615. In bladder, due to fewer patches available in the small slide, only seven salient and seven nonsalient examples are drawn from the second patient. Average AUROC was calculated for ten such draws, achieving 0.7929±0.1109 and 95% CI of 0.7176-0.8763. Over all 7 salient and 17 nonsalient patches used from the second bladder patient, the AUROC is 0.7437. These nonoverlapping confidence intervals are evidence the bladder cancer classifier distinguishes salient from nonsalient patches less well than the prostate cancer classifier, and a Mann-Whitney U test indeed finds the difference in classifier performance by these ten draws each from bladder and prostate is significant (p=0.0001325) (Fig 9).

**Figure 10:**
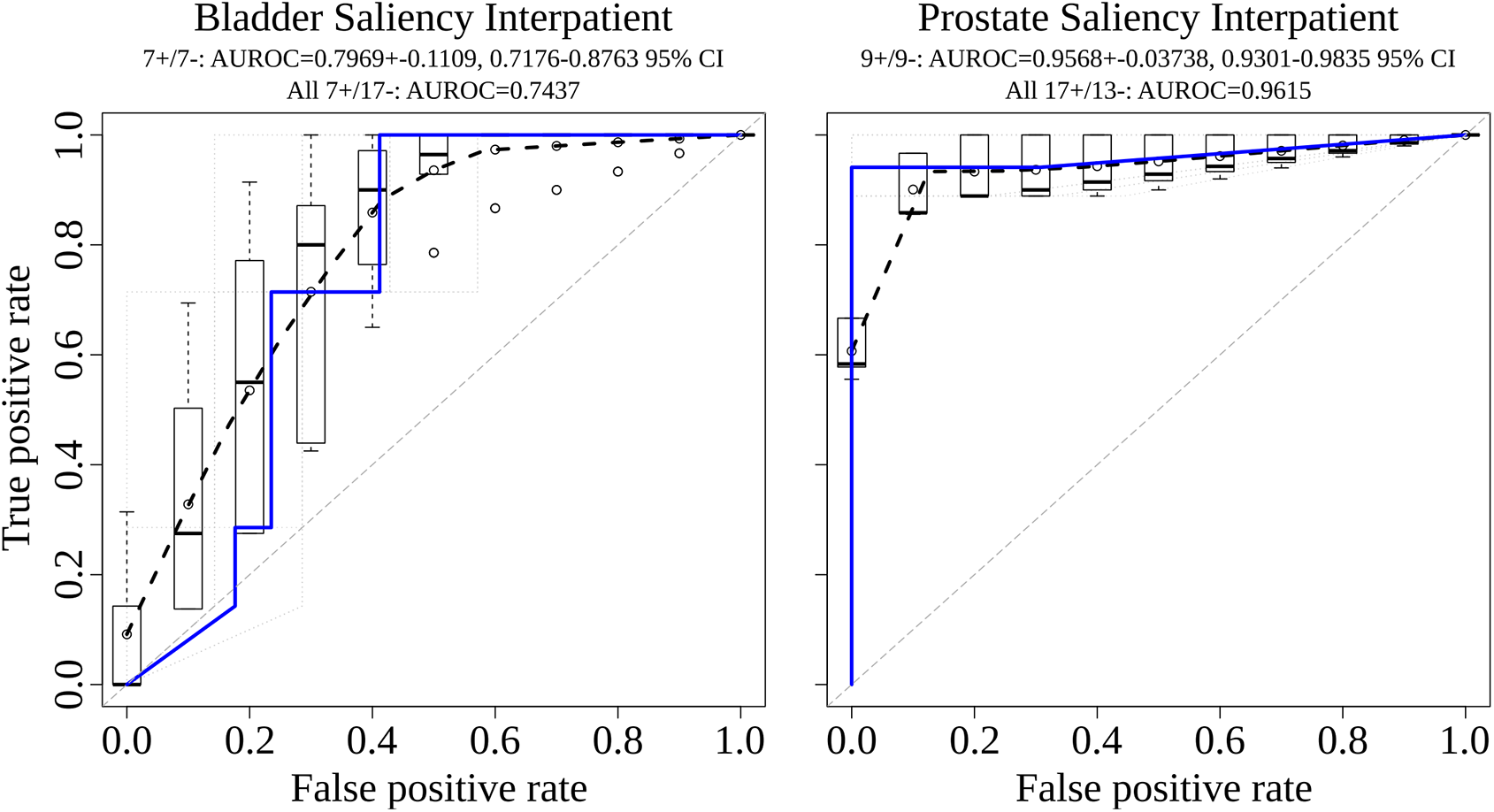
Interpatient area under the receiver operating characteristic [AUROC] for bladder and prostate, with dashed black curve for average AUROC over draws of the data and blue line for all data used from the patient.

The deep convolutional network CaffeNet emits a score from 0 to 1 when predicting if an image patch is salient or not, and when taking a score of greater than 0.5 to be salient, the p-value from Fisher’s Exact Test is 1.167e-7 in prostate (16 true positives, 1 false negative, 0 false positives, 13 true negatives) and 0.009916 in bladder (7 true positives, 0 false negatives, 7 false positives, 10 true negatives), indicating our trained CaffeNet classifier accurately predicts salient and nonsalient regions in both tissues.

## 5 Conclusion

Collecting image-based expert annotations for the deluge of medical data at modern hospitals is one of the biggest bottlenecks for the application of large-scale supervised machine learning. We address this with a novel framework that combines a commodity camera, 3D-printed mount, and software stack to build a predictive model for saliency on whole slides, i.e. where a pathologist looks to make a diagnosis. The registered regions from the digital slide scan are markedly higher quality than the camera frames, since they do not suffer from debris, vignetting, and other artifacts. The proposed CNN is able to predict salient slide regions with a test accuracy of 85-91%. We plan to scale up this pilot study to more patients, tissues, and pathologists.

**Table 1:** Accuracies of ten trials of three-fold cross validation in bladder. Validation and test accuracies for a single slide video of urothelial carcinoma (patient 1, slide at upper left in Fig 2, performance plotted at left in Fig 9)), left side of the slide versus right side. Mean test error training/validating on left and testing on right [leftright] is 0.781 at stdev 0.0423 with 95% confidence interval from 0.750 to 0.811 (df=9, Student’s t), while mean test error right to left [rightleft] is 0.922 at stdev 0.0468 with 95% confidence interval from 0.889 to 0.956 (df=9, Student’s T). In cases where best validation accuracies (highlighted in yellow) tie in multiple models, all tied models are used to evaluate test accuracy and their results averaged. Testing the best classifier (highlighted in cyan, highest test accuracy on this and other folds, secondarily highest mean validation accuracy) on draws of the data on the second bladder patient, accuracies are 0.643, 0.786, 0.714, 0.786, 0.714, 0.714, 0.643, 0.571, 0.643, and 0.571.

| Direction | Trial | Fold0<br>Valid | Fold1<br>Valid | Fold2<br>Valid | Valid<br>Acc | Fold0<br>Test | Fold1<br>Test | Fold2<br>Test | Test<br>Acc |
| --- | --- | --- | --- | --- | --- | --- | --- | --- | --- |
| leftright | 0 | 0.9466 | 0.5850 | 0.9680 | 0.8332 | 0.8333 | 0.7222 | 0.7778 | 0.7778 |
| leftright | 1 | 0.8852 | 0.9070 | 0.9070 | 0.8997 | 0.7778 | 0.7222 | 0.7778 | 0.7500 |
| leftright | 2 | 0.9218 | 0.8602 | 0.8832 | 0.8884 | 0.8333 | 0.7222 | 0.7778 | 0.8333 |
| leftright | 3 | 0.7640 | 0.7812 | 0.7120 | 0.7524 | 0.7778 | 0.7778 | 0.7778 | 0.7778 |
| leftright | 4 | 0.7590 | 0.6576 | 0.5134 | 0.6433 | 0.8333 | 0.7778 | 0.7778 | 0.8333 |
| leftright | 5 | 0.9268 | 0.7416 | 0.5088 | 0.7257 | 0.8333 | 0.7778 | 0.7778 | 0.8333 |
| leftright | 6 | 0.8028 | 0.7988 | 0.7048 | 0.7688 | 0.7222 | 0.7778 | 0.7778 | 0.7222 |
| leftright | 7 | 0.7318 | 0.8402 | 0.9088 | 0.8269 | 0.7778 | 0.7778 | 0.7778 | 0.7778 |
| leftright | 8 | 0.9572 | 0.7608 | 0.8418 | 0.8533 | 0.7778 | 0.8333 | 0.8333 | 0.7778 |
| leftright | 9 | 0.7492 | 0.8774 | 0.9860 | 0.8709 | 0.7778 | 0.7778 | 0.7222 | 0.7222 |
| rightleft | 0 | 0.8802 | 0.8528 | 0.8554 | 0.8628 | 1.0000 | 0.8889 | 0.9444 | 1.0000 |
| rightleft | 1 | 0.7662 | 0.5982 | 0.9364 | 0.7669 | 0.8889 | 0.9444 | 1.0000 | 1.0000 |
| rightleft | 2 | 0.9492 | 0.8560 | 0.7308 | 0.8453 | 0.9444 | 0.8889 | 0.9444 | 0.9444 |
| rightleft | 3 | 0.5404 | 0.8206 | 0.8368 | 0.7326 | 0.9444 | 0.9444 | 0.8889 | 0.8889 |
| rightleft | 4 | 0.6560 | 0.7114 | 0.6748 | 0.6807 | 0.8889 | 0.8889 | 0.8333 | 0.8889 |
| rightleft | 5 | 0.8932 | 0.7062 | 0.7310 | 0.7768 | 0.9444 | 0.8889 | 0.8333 | 0.9444 |
| rightleft | 6 | 0.8560 | 0.8540 | 0.9966 | 0.9022 | 0.8889 | 0.9444 | 0.8889 | 0.8889 |
| rightleft | 7 | 0.8362 | 0.8560 | 0.7978 | 0.8300 | 0.8333 | 0.8889 | 1.0000 | 0.8889 |
| rightleft | 8 | 0.7200 | 0.8546 | 0.9740 | 0.8495 | 1.0000 | 0.9444 | 0.8889 | 0.8889 |
| rightleft | 9 | 0.8634 | 0.8634 | 0.6904 | 0.8057 | 0.8333 | 0.9444 | 1.0000 | 0.8889 |

**Table 2:** Accuracies of ten trials of three-fold cross validation in prostate. Validation and test accuracies for a single slide video of prostate adenocarcinoma (patient 1, slide at upper right in Fig 2, performance plotted at right in Fig 9)), left side of the slide versus right side. Mean test error training/validating leftright is 0.781 at stdev 0.0423 with 95% confidence interval from 0.750 to 0.811 (df=9, Student’s t), while mean test error rightleft is 0.922 at stdev 0.0468 with 95% confidence interval from 0.889 to 0.956 (df=9, Student’s T). Testing the best classifier on draws of the data on the second prostate patient, accuracies are 0.944, 1, 0.944, 0.944, 1, 0.944, 0.944, 0.944, 1, and 1.

| Direction | Trial | Fold0<br>Valid | Fold1<br>Valid | Fold2<br>Valid | Valid<br>Acc | Fold0<br>Test | Fold1<br>Test | Fold2<br>Test | Test<br>Acc |
| --- | --- | --- | --- | --- | --- | --- | --- | --- | --- |
| leftright | 0 | 0.9992 | 0.9946 | 1.0000 | 0.9979 | 0.7778 | 0.7778 | 0.7778 | 0.7778 |
| leftright | 1 | 0.9512 | 0.7282 | 0.9994 | 0.8929 | 0.8889 | 0.8889 | 0.8889 | 0.8889 |
| leftright | 2 | 0.9550 | 1.0000 | 0.6530 | 0.8693 | 0.8889 | 0.7778 | 0.7222 | 0.7778 |
| leftright | 3 | 0.8636 | 0.9992 | 1.0000 | 0.9543 | 0.8889 | 0.7778 | 0.9444 | 0.9444 |
| leftright | 4 | 0.8276 | 0.7760 | 0.9940 | 0.8659 | 0.8889 | 0.7778 | 0.8889 | 0.8889 |
| leftright | 5 | 0.8654 | 0.9986 | 1.0000 | 0.9547 | 0.9444 | 0.9444 | 0.9444 | 0.9444 |
| leftright | 6 | 0.8560 | 0.9862 | 0.9992 | 0.9471 | 0.8889 | 0.8889 | 0.8889 | 0.8889 |
| leftright | 7 | 0.8674 | 1.0000 | 0.9984 | 0.9553 | 0.8889 | 0.8333 | 0.8889 | 0.8333 |
| leftright | 8 | 0.9560 | 0.8560 | 0.8760 | 0.8960 | 0.8889 | 0.8333 | 0.9444 | 0.8889 |
| leftright | 9 | 0.6846 | 0.9992 | 0.9560 | 0.8799 | 0.8333 | 0.8333 | 1.0000 | 0.8333 |
| rightleft | 0 | 0.9786 | 0.7760 | 0.9146 | 0.8897 | 0.8889 | 0.9444 | 1.0000 | 0.8889 |
| rightleft | 1 | 1.0000 | 0.7292 | 0.8460 | 0.8584 | 1.0000 | 0.9444 | 1.0000 | 1.0000 |
| rightleft | 2 | 0.7130 | 0.9512 | 0.8676 | 0.8439 | 0.8889 | 1.0000 | 0.8333 | 1.0000 |
| rightleft | 3 | 0.9998 | 1.0000 | 0.9664 | 0.9887 | 0.9444 | 0.9444 | 1.0000 | 0.9444 |
| rightleft | 4 | 0.7760 | 1.0000 | 0.8842 | 0.8867 | 0.8889 | 0.9444 | 1.0000 | 0.9444 |
| rightleft | 5 | 0.9758 | 0.9984 | 0.5926 | 0.8556 | 0.9444 | 0.8889 | 0.9444 | 0.8889 |
| rightleft | 6 | 0.6344 | 0.9770 | 1.0000 | 0.8705 | 0.8889 | 1.0000 | 1.0000 | 1.0000 |
| rightleft | 7 | 0.7760 | 0.9028 | 1.0000 | 0.8929 | 0.8889 | 1.0000 | 1.0000 | 1.0000 |
| rightleft | 8 | 0.8560 | 0.8560 | 0.9992 | 0.9037 | 0.9444 | 0.9444 | 0.9444 | 0.9444 |
| rightleft | 9 | 0.8560 | 0.9412 | 0.8538 | 0.8837 | 1.0000 | 1.0000 | 1.0000 | 1.0000 |

## Acknowledgments

AJS was supported by NIH/NCI grant F31CA214029 and the Tri-Institutional Training Program in Computational Biology and Medicine (via NIH training grant T32GM083937). This research was funded in part through the NIH/NCI Cancer Center Support Grant P30CA008748. AJS thanks Terrie Wheeler, Du Cheng, and the Medical Student Executive Committee of Weill Cornell Medical College for free 3D printing access, instruction, and support. AJS thanks Mariam Aly for taking the photo of the camera on the orange 3D-printed mount in Fig 1, and attention discussion. We acknowledge fair use of part of a doctor stick figure image in Fig 1 from 123rf.com. AJS thanks Mark Rubin for helpful pathology discussion. AJS thanks Paul Tatarsky and Juan Perin for Caffe install help on the Memorial Sloan Kettering supercomputer. Several GPUs were used in this research, one of which we gratefully acknowledge was provided by NVIDIA Corporation as part of a GPU Research Center award to TJF.

## Footnotes

1 ImageJ SURF is released under the GNU GPL and is available for download from http://labun.com/imagej-surf/

